# Prevention of mRNA vaccine-induced anaphylaxis by peripheral cyclooxygenase inhibitors in an anti-PEG hyperimmune pig model: clinical relevance for nanomedicine-induced infusion reactions

**DOI:** 10.64898/2026.05.27.727997

**Authors:** Bálint András Barta, Tamás Radovits, Attila Balázs Dobos, Sylvia Spiesshofer, Ákos Gergely Tóth, Gleb Kornev, Alberto Gabizon, Béla Merkely, János Szebeni

**Affiliations:** Heart and Vascular Center, Semmelweis University, Budapest, Hungary; Helmsley Cancer Center, Shaare Zedek Medical Center, Jerusalem, Israel; Nanomedicine Research and Education Center, Department of Clinical Pathophysiology, Semmelweis University, Budapest, Hungary; SeroScience LCC, Budapest, Hungary

## Abstract

Anti-polyethylene glycol (PEG) hyperimmune pigs, immunized against PEG, provide a sensitive experimental model for the rare anaphylactic reactions induced by mRNA-PEGylated lipid nanoparticle (LNP)-based COVID-19 vaccines, such as Comirnaty. These pseudo-allergic infusion reactions can usually be prevented or attenuated by multicomponent anti-inflammatory premedication regimens; however, no established protocol exists for mRNA-LNP-based COVID-19 vaccines. The aim of the present study was to identify an effective premedication strategy capable of preventing or attenuating these reactions in hypersensitive subjects, using the hyperimmune porcine model. We compared the protective effects of individual pretreatment components; dexamethasone, famotidine, levocetirizine, acetaminophen, diclofenac, indomethacin, by analyzing hemodynamic endpoints (systemic and pulmonary arterial pressure, pulse pressure). All tested compounds modulated Comirnaty-induced anaphylactic responses; however, only cyclooxygenase (COX) inhibitors provided complete protection against anaphylaxis and other abnormal processes. This finding is consistent with the low incidence of infusion reactions to cancer nanomedicines at the Shaare Zedek Oncology Center in Israel which uses COX-inhibitors as premedication. Given that most currently used human infusion-reaction prevention protocols do not include COX inhibitors, and that steroid-containing regimens may potentially counteract vaccine efficacy, our results suggest that COX inhibitors may offer a clinically effective standalone option or form the basis of simplified premedication regimens for preventing this life-threatening condition.

## INTRODUCTION

Anaphylaxis, a life-threatening hypersensitivity reaction (HSR), leading the safety warnings of Pfizer-BioNTech’ mRNA-lipid nanoparticle (LNP)-based vaccine (Comirnaty) [1] was a relatively rare adverse event (AE) that occurred in a few cases in thousands of vaccine recipients during the height of vaccination campaign against COVID-19 [2–7] Since then, the use of these vaccines has declined, yet there are several reasons warranting sustained attention to, and systematic investigation of vaccine-induced anaphylaxis. These include the continued use of mRNA vaccines all over the world with FDA/CDC recommendation in the USA only for selected elderly populations; the 40-fold excess of incidence rate over that reported for seasonal influenza vaccines; the ongoing expansion of mRNA-LNP technology to indications beyond COVID-19 in the face of increasing percentage of severely allergic individuals in the society (up to ∼2% of the population) in whom the anaphylaxis risk is several-fold higher than in the healthy population [8]. Collectively, these considerations underscore that the prediction and prevention of anaphylaxis by RNA-LNP vaccines and therapeutics represent a major unmet medical need.

Regarding the mechanism of vaccine-induced HSRs and anaphylaxis, although the clinical manifestations, such as the acute tachycardia, dyspnea, tachypnea, apnea, hypo- or hypertension, generalized or localized flushing or rash, and angioedema are consistent with type I allergy, the pre-sensitization with a vaccine component, yielding specific IgE antibodies, can explain only a negligible fraction of vaccine-induced reactions [9]. The overwhelming majority of reactions represent nonspecific innate immune response involving complement (C) activation. Although the pathomechanism is far more complex than solely anaphylatoxin action, the pivotal role of C activation in the phenomenon led to the mnemonic term, complement activation-related pseudoallergy/pseudoanaphylaxis (CARPA).

CARPA is a well-recognized, ubiquitous consequence of intravenously administered reactogenic nanoparticles, including liposomes, micellar solvents, contrast media, and complex biologicals [10–12]. Its involvement in vaccine-induced HSRs is a natural consequence of the nanoparticle-based composition of these vaccines and was demonstrated in porcine studies in which the vaccine was administered intravenously [13]. As an innovative breakthrough regarding the pig model, we reported [7] that pre-immunization of the animals with a placebo, drug-free, PEGylated nanoparticle (Doxebo) sensitized them for anaphylaxis induction by PEGylated nanoparticles, leading Barta et al. to establish a uniquely sensitive and reproducible porcine model of Comirnaty-induced anaphylaxis [14]. Beyond the practical advances, the model provided evidence for a causal relationship between anti-PEG antibody-mediated C activation and anaphylactoid reactions to Comirnaty [14].

If vaccine-induced HSRs and anaphylaxis share the CARPA mechanism known from nanomedicine-induced HSRs [15], it is reasonable to ask whether vaccine-induced reactions might also be attenuated or prevented by the standard premedication regimens applied for decades against drug-induced HSRs, commonly known as infusion reactions. Such premedication protocols typically involve combinations of anti-inflammatory agents and antihistamines with schedules and regiments variably applied in different institutions for different drugs (Table 1). Nevertheless, as seen in Table 1, neither of these “standard-of-care” regimens provide full protection against HSRs/anaphylaxis, with reported incidences remaining in the 5-20% range. One exception is the Shaare Zedek protocol [16], used in the human studies presented here, which includes COX-inhibitors leading to substantial reduction of reaction risk, as presented below.

**Table 1.** Reactogenic drugs, premedication methods and HSR incidence.

| Reactogenic drug | Premedication regimen | Additional strategies for reaction prevention | HSR (% incidence) | References |
| --- | --- | --- | --- | --- |
| Doxil<sup>®</sup>, Caelix<sup>®</sup><br>(pegylated liposomal doxorubicin) | • Dexamethasone<br>• Diphenhydramine<br>• Ranitidine<br>• Famotidine<br>• Acetaminophen | • slow infusion with stepwise rate escalation<br>• tachyphylaxis induction with placebo liposomes<br>• close monitoring for first 20 minutes<br>• emergency medications at bedside<br>• adjust infusion rate if reaction occurs<br>• close monitoring | 7-40 | [17-20] |
| AmBisome<sup>®</sup><br>(liposomal amphotericin B) | • Dexamethasone<br>• Diphenhydramine<br>• Ranitidine<br>• Famotidine<br>• Acetaminophen<br>• Meperidine | • slow infusion $\geq 2$ h<br>• apply test | 5-50 | [21-23] |
| Promitil<sup>®*</sup><br>(liposomal mitoxantron) | • *Dexamethasone*<br>• *Diphenhydramine*<br>• *Ranitidine*<br>• *Acetaminophen*<br>• *COX inhibitors (ibuprofen, diclofenac, indomethacin)* | • *slow infusion*<br>• *close monitoring* | 10-20 | [24, 25] |
| Onpattro<sup>®</sup> (patisiran, siRNA-LNP) | • Dexamethasone<br>• Diphenhydramine<br>• Ranitidine<br>• Famotidine<br>• Acetaminophen | • 80 min infusion adjusted as needed | 18-20 | [26-28] |
| Marqibo<sup>®</sup><br>(liposomal vincristine sulfate) | • Dexamethasone<br>• Diphenhydramine<br>• Ranitidine<br>• Famotidine<br>• Acetaminophen | • slow (10-15 min) infusion | 5-15 | [29-31] |
| Onivyde<br>(liposomal irinotecan) | • Diphenhydramine<br>• corticosteroid | • 90 min infusion<br>• close monitoring | 3-10 | [32-35] |
| Lipusu –<br>(liposomal paclitaxel) | • Dexamethasone<br>• antiemetics | • 1-2 h infusion | 10-30 | [36-38] |
| Lipoplatin<br>(liposomal platinum) | • Dexamethasone<br>• Diphenhydramine<br>• Ranitidine<br>• antiemetics | • Slow infusion. | 5-10 | [39, 40] |
| Abelcet<br>(liposomal doxorubicin) | • Acetaminophen<br>• antihistamines | • 2-4 h infusion<br>• close monitoring | 15-40 | [41-43] |

\*Shaare Zedek protocol (italicized)

Based on the above gaps in safety, the primary objective of the present study was to use the anti-PEG hyperimmune porcine model of Comirnaty-induced anaphylaxis [14] to evaluate the efficacy of different premedication protocols in preventing vaccine-induced reactions, dissecting the individual contributions of selected components. To align the pig data with human experience, we also analyzed retrospectively the reaction incidence and severity in cancer patients treated with Doxil and Promitil after applying the premedication protocol specified in the text.

## METHODS

### Animals

Mixed-breed Yorkshire/Hungarian White Landrace pigs of both sexes (2-3 months old, 20-28 kg) were obtained from the Animal Breeding, Nutrition and Meat Science Research Institute, Hungarian University of Agriculture and Life Sciences (Herceghalom, Hungary).

### Ethics

The investigation conformed to the EU Directive 2010/63/EU and the Guide for the Care and Use of Laboratory Animals used by the US National Institutes of Health (NIH Publication No.85-23, revised 1996). The experiments were approved by the Ethical Committee of Hungary for Animal Experimentation (permission number: PE/EA/843-7/2020.

### Drugs

The drugs used for premedication were obtained from the following sources: acetaminophen, PARAMAX Rapid 500 mg (Vitabalans Oy, Hämeenlinna, Finland); famotidine, Quamatel (Gedeon Richter Plc., Budapest, Hungary); dexamethasone, KRKA (Novo mesto, Slovenia); levocetirizine, Novocetrin (Teva Gyógyszergyár Zrt., Debrecen, Hungary); diclofenac-ratiopharm (Teva Gyógyszergyár Zrt., Debrecen, Hungary); and indomethacin (Sigma-Aldrich/Merck KGaA, Darmstadt, Germany).

### Preparation of Doxebo

The freeze-dried lipid components of Doxil were hydrated in 10 mL sterile pyrogen-free normal saline by vortexing for 2-3 min at 70^◦^C to form multilamellar vesicles (MLVs). The MLVs were downsized through 0.4 and 0.1 μm polycarbonate filters in two steps, 10 times each, using a 10 mL extruder barrel from Northern Lipids (Vancouver, British Columbia, Canada) at 62 ^◦^C. Liposomes were suspended in 0.15 M NaCl/ 10 mM histidine buffer (pH 6.5). The size distribution (Z-average): 81.17 nm and phospholipid concentration (12.6 mg/mL) were determined.

### Sensitization of pigs against PEGylated liposomes by preimmunization with Doxebo

Baseline (“pre-immune”) blood samples were taken from the pigs followed by immunization by way of infusion of 0.1 mg PL/kg Doxebo via the ear vein (suspended in 20 mL of saline) at a speed of 1 mL/min. The animals were then placed back into their cages until the 2nd blood sampling 9-10 days later, to screen for anti-PEG Ab induction. The animals were then subjected to Comirnaty challenges and CARPA assay within 2 weeks.

### Experimental protocol

Pigs were sedated with Ketamin/Xilazine, anesthetized with isoflurane (2-3 % in O2) and intubated with endotracheal tubes to maintain free airways and to enable controlled ventilation if spontaneous breathing stopped during the experiment. After iodine (10 %, povidone) disinfection of the skin, the pigs were subjected to surgery to insert various catheters into their circulation, namely: (a) a Swan-Ganz catheter (Arrow AI-07124 single- lumen balloon wedge pressure catheter 5 Fr., 110 cm, Arrow International Inc, Reading, PA, USA), into the pulmonary artery via the right external jugular vein (in order to measure the pulmonary arterial pressure (PAP); (b) the left femoral artery to record the systemic arterial pressure (SAP); (c) the left external jugular vein for saline and drug infusion; (d) into the left femoral vein for blood sampling.

The above hemodynamic were recorded by a physiological monitoring system (Pulsion Medical Systems SE, Munich, Germany) and Powerlab (ADInstruments, Bella Vista, Australia). End-tidal pCO2, O2 saturation, ventilation rate and body temperature were also continuously measured. At the end of the experiments the animals were sacrificed with pentobarbital (120 mg/kg i.v.) and concentrated potassium chloride.

Continuous physiologic data were aligned to the time of challenge, and the minute immediately preceding challenge was used as baseline. Hemodynamic variables were summarized at 1-minute resolution, whereas respiratory and hematologic variables were available at predefined sparse post-challenge time points. For all analyses, only animals belonging to the predefined pretreatment groups were included. To account for baseline differences between animals, outcomes were expressed both as absolute change from baseline and as percentage change from baseline. Hemodynamic analyses focused on the early post-challenge period (10 minutes).

### Comparison of Comirnaty-induced anaphylaxis following different premedications

After 15-30 min adaptation, hyperimmune pigs were subjected to the following premedications; 1) the corticosteroid dexamethasone (4 mg p.o.); 2) the analgesic and antipyretic drug acetaminophen (500 mg p.o.), 3) a combination of antihistamines comprising of the H1-receptor antagonist, levocetirizine (5 mg p.o.) and H2-receptor antagonist, famotidine (40 mg p.o.), 4) the same antihistamine combination supplemented with the COX inhibitor diclofenac (100 mg p.o.), (5) diclofenac alone (100 mg p.o.), or 6) another COX inhibitor indomethacin alone (100 mg i.v.). 1.5 hours later baseline blood samples were taken, and the animals were first injected i.v. with 5 mL PBS, as negative control, then with 1/3 human vaccine dose (HVD) of Comirnaty containing 10 μg mRNA, corresponding to 0.4-0.5 μg/kg mRNA. Finally 0.1 mg/kg zymosan was injected i.v. as positive control. The un-premedicated pigs were treated similarly, without giving the premedication drug combination.

Hemodynamic and cardiopulmonary monitoring, detailed above, was continuous after vaccine challenge during the primary reaction phase, up to 20-30 min. The readings were averaged at one-minute intervals and are presented as bars ± SEM. Blood withdrawal for blood cell and inflammatory biomarker changes was performed at the times indicated for each experiment. The analyzed hemodynamic endpoints were mean pulmonary arterial pressure (mPAP), mean systemic arterial pressure (mSAP), as well as corresponding pulse amplitude (pSAP).

### Schematic overview of study design

**Figure 1** illustrates the experimental stages and timeline of the study, extending our previous report on Comirnaty-induced anaphylaxis in anti-PEG hyperimmune pigs [1]. The diagram summarizes the sequence of animal sensitization, premedication regimens, intravenous Comirnaty challenge, real-time hemodynamic monitoring, blood sampling, and subsequent laboratory analyses.

**Figure 1.**
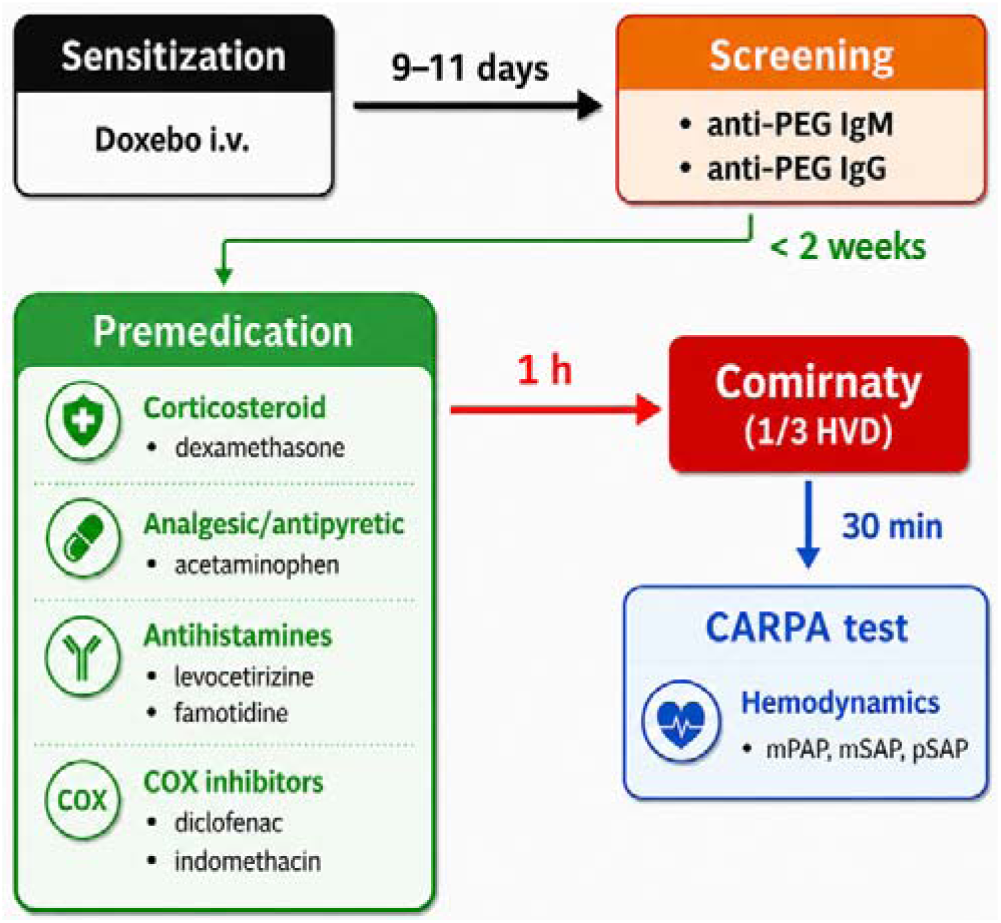
Experimental workflow and timeline of the present study. The sequence highlights the key stages of procedures, their scheduling and duration.

### Statistical analysis of hemodynamic, respiratory, and hematologic responses

The main analysis tested whether pretreatment changed the responses after Comirnaty injection. Separate statistical models were used for each measured variable. These models compared the treatment groups over time and accounted for repeated measurements from the same animal. The responses were adjusted to each animal’s baseline value. We tested whether the groups differed overall and whether their time courses were different. Group differences were also compared at selected time points after injection. To quantitate each response, we calculated the area under the effect curve (AUEC) which captures the overall magnitude of deviation from baseline irrespective of direction. The absolute AUEC was used to measure the total change from baseline, and group differences in AUEC were then compared between untreated and pretreatment groups. P values from multiple pairwise comparisons were adjusted using the Benjamini-Hochberg procedure. All tests were two-sided, and adjusted p values were used for post hoc inference.

## RESULTS

### Doxebo Sensitization of Pigs for Anaphylactic Reactions to Comirnaty

Although i.v. administration of Doxebo, a PEGylated liposome, is technically considered treatment rather than immunization, we refer to this process as immunization because of the strong immunogenic response illustrated in Supplementary SFig. 1. In the present study, the anti-PEG IgM and anti-PEG IgG increased in 9-10 days approximately 300X and 17X fold, respectively.

### Comirnaty-induced physiological changes in anti-PEG hyperimmune pigs without premedication

Figure 2 illustrates the physiological changes of CARPA-characteristic biomarkers following administration of 1/3 HVD of Comirnaty to anti-PEG hyperimmune pigs without any premedication. In Panels A and B, a representative reaction is shown. After a stable saline baseline with no detectable changes in pulmonary arterial pressure (PAP; A) or systemic arterial pressure (SAP; B), i.v. administration of the vaccine induced an abrupt and profound hemodynamic disturbance within 1-2 minutes after the start of bolus injection. PAP rose steeply (A) and developed large-amplitude oscillations, reflecting circulatory abnormality. Almost simultaneously, SAP began to fall rapidly (B) and reached shock-level hypotension within a few minutes, reflecting sudden impairment of left-ventricular filling secondary to pulmonary vasoconstriction. At the nadir of systemic hypotension, cardiopulmonary resuscitation (CPR) was initiated, resulting in a prompt recovery and transient overshoot of SAP (Fig 2B), while PAP gradually declined but remained transiently elevated compared with baseline (Fig 2A). After hemodynamic stabilization, administration of zymosan (Zym), used as a positive control for complement (Fig 2 A, B) activation. Panels C and D show the mean ± SEM changes in PAP and SAP, respectively, obtained from 10 pigs. The relatively small SEM values during the rising and falling phases of both pressures demonstrate that the timing and magnitude of these hemodynamic changes were highly consistent across animals. Panels E and F show the paralleling changes in blood inflammatory markers, the anaphylatoxin C3a and thromboxane A2 metabolite, TXB2, which are pivotal biomarkers of humoral and cellular innate immune stimulation involving C activation and downstream eicosanoid release by secretory immune cells. The close coupling of the timing of these processes together with the phenotypic similarity between Comirnaty- and zymosan-induced responses provide strong support for the conclusion that the vaccine reactions are due, at least in part, to C activation-related pseudoallergy (CARPA) [1].

**Figure 2.**
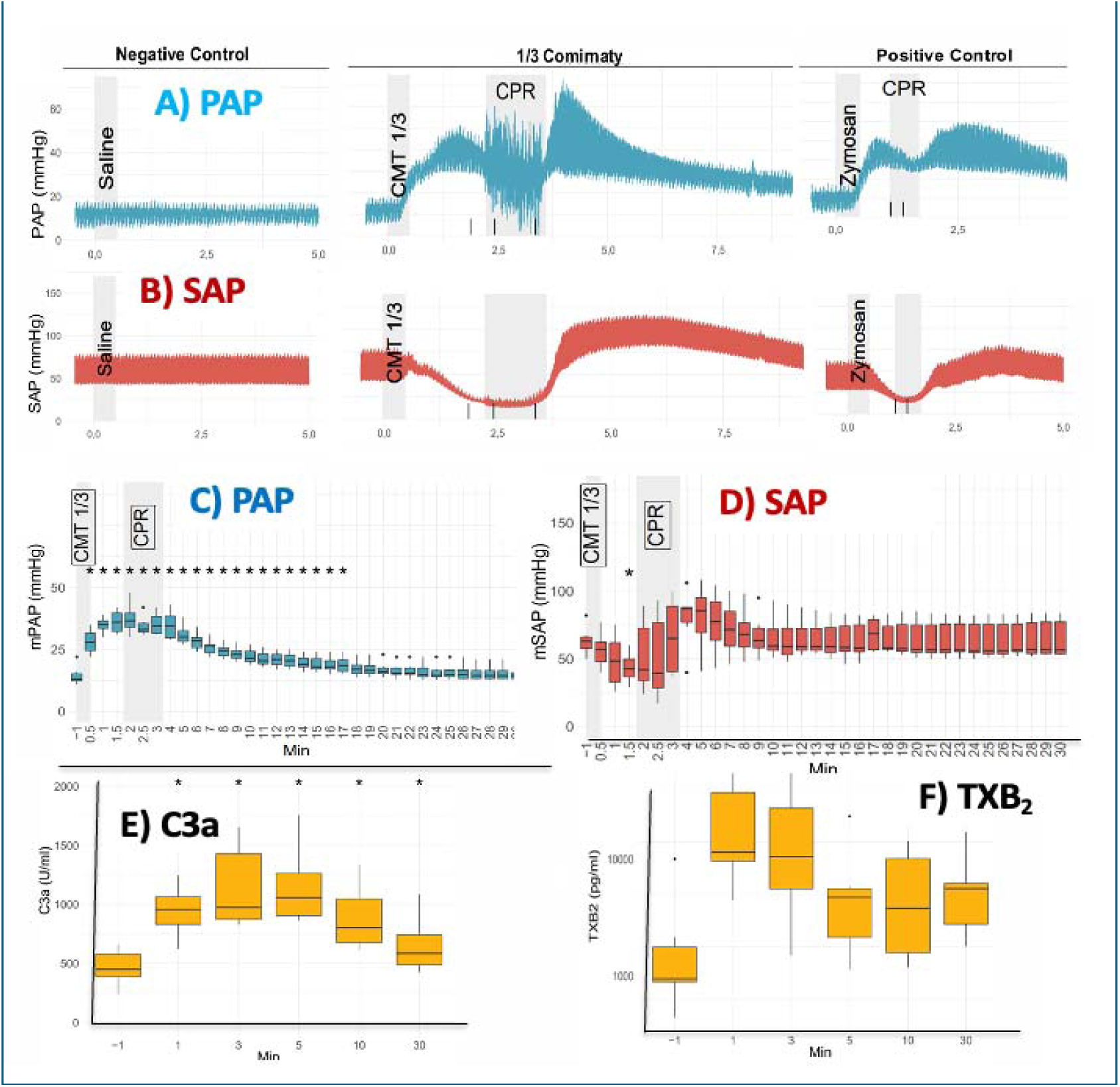
Real-time monitoring of the hemodynamic changes and paralleling systemic inflammatory response caused by Comirnaty in anti-PEG hyperimmune pigs. Continuous recordings of pulmonary arterial pressure (PAP; A) and systemic arterial pressure (SAP; B) during the three phases of the experiment: (1) baseline saline infusion; (2) injection of 1/3 of the human dose of Comirnaty (HVD) and (3) injection of zymosan, the gold standard of C and innate immune cell activation. CPR, cardiopulmonary resuscitation. Panels C and D show mean ± SEM PAP and SAP responses, respectively, following CMT injection (n = 10). Panels E and F show the plasma C3a and TXB□ changes, respectively, during the anaphylactic reactions. Reproduced from [14] with permission.

### Effects of premedication regimens on Comirnaty-induced pulmonary hypertension

Figure 3A shows the effects of Comirnaty on mean PAP in premedicated and control un-premedicated pigs with the treatments specified in the individual panels. To eliminate baseline variability, PAP responses are expressed as the percentage increase in PAP relative to the mean baseline value. In each of 10 untreated pigs the PAP response curves display the robust pulmonary hypertension detailed in Fig. 3A. The pretreatment with dexamethasone and antihistamines failed to provide any protection against the reaction, while acetaminophen suppressed the reaction in 2 out of 3 pigs. In sharp contrast, the antihistamine/diclofenac combination as well as diclofenac and indomethacin, both applied alone, produced essentially complete inhibition of the rise of PAP in all of 18 pigs injected with the vaccine. Figure 3B shows the 95% confidence bands for these changes. The broad trajectories overlap among the non-premedicated reactor group and the groups premedicated with dexamethasone, antihistamines and acetaminophen, indicating no significant differences. In contrast, the confidence-band trajectories of animals premedicated with COX inhibitors are narrow, remain close to baseline, and are sharply separated from the above groups, consistent with significant stabilization of PAP.

**Figure 3.**
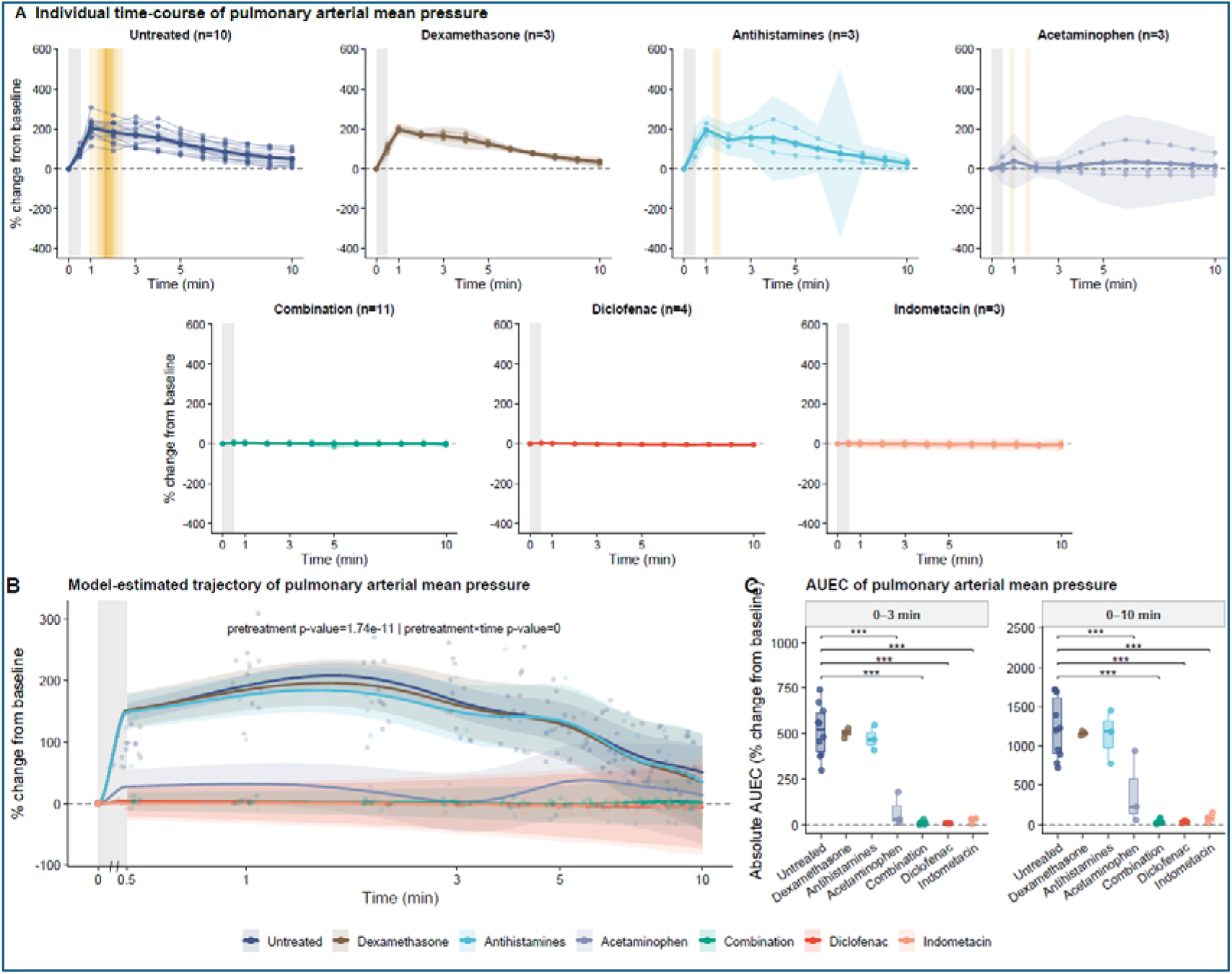
Effects of premedication regimens on Comirnaty-induced changes in mean PAP in anti-PEG hyperimmune pigs, expressed as percentage change from baseline. **(A)** PAP responses to CMT in pigs without premedication and after premedication with dexamethasone (4mg p.o.) acetaminophen (500 mg p.o.), the combination of levocetirizine (5 mg p.o.) plus famotidine (40 mg p.o.), the same antihistamine combination supplemented with diclofenac (100 mg p.o.), diclofenac alone (100 mg p.o.), or indomethacin alone (100 mg i.v.). Thin lines represent mean PAP values in individual animals, thick lines indicate group means, with animal numbers shown in brackets, and shaded areas showing the 95% confidence bands. The yellow vertical band indicates resuscitation with noradrenaline, where required. **(B)** Superimposed PAP response curves with their 95% confidence bands over time. The shaded vertical intervals in A and B mark the early post-injection response window. **(C)** Area under the effect curve (AUEC) calculated for the early response phase, 0-3 min, and for the full observation period, 0-10 min.

Panel C quantifies all these differences among treatment groups relative to the non-premedicated control, based on the area under the curve within the dynamic range of the hemodynamic response. These data statistically confirm the information in panels A and B, namely that both COX inhibitors exerted highly significant and essentially complete inhibition of Comirnaty-induced pulmonary hypertension.

### Effects of different premedication regimens on Comirnaty-induced systemic blood pressure changes

Figure 4A shows the effect of Comirnaty on mean SAP in the same experiments presented in Fig. 4A-C. In contrast to the robust and uniform PAP elevation, the averaged systemic arterial pressure response appeared more variable and showed only limited net deviation from baseline. However, this should not be interpreted as absence of systemic reactogenicity. In animals showing a clear reaction, Comirnaty predominantly induced a marked fall in SAP, in some cases reaching life-threatening severity. To avoid early termination of the experiments and to maintain systemic perfusion, intravenous catecholamines had to be administered in the most severely affected pigs. These rescue interventions inevitably modified the subsequent SAP course, contributing to the apparent biphasic character of the averaged curves and to the broad 95% confidence bands. Accordingly, the early 0-3 min AUEC interval is particularly informative, because it best reflects the vaccine-induced hemodynamic response before catecholamine effects became dominant.

**Figure 4.**
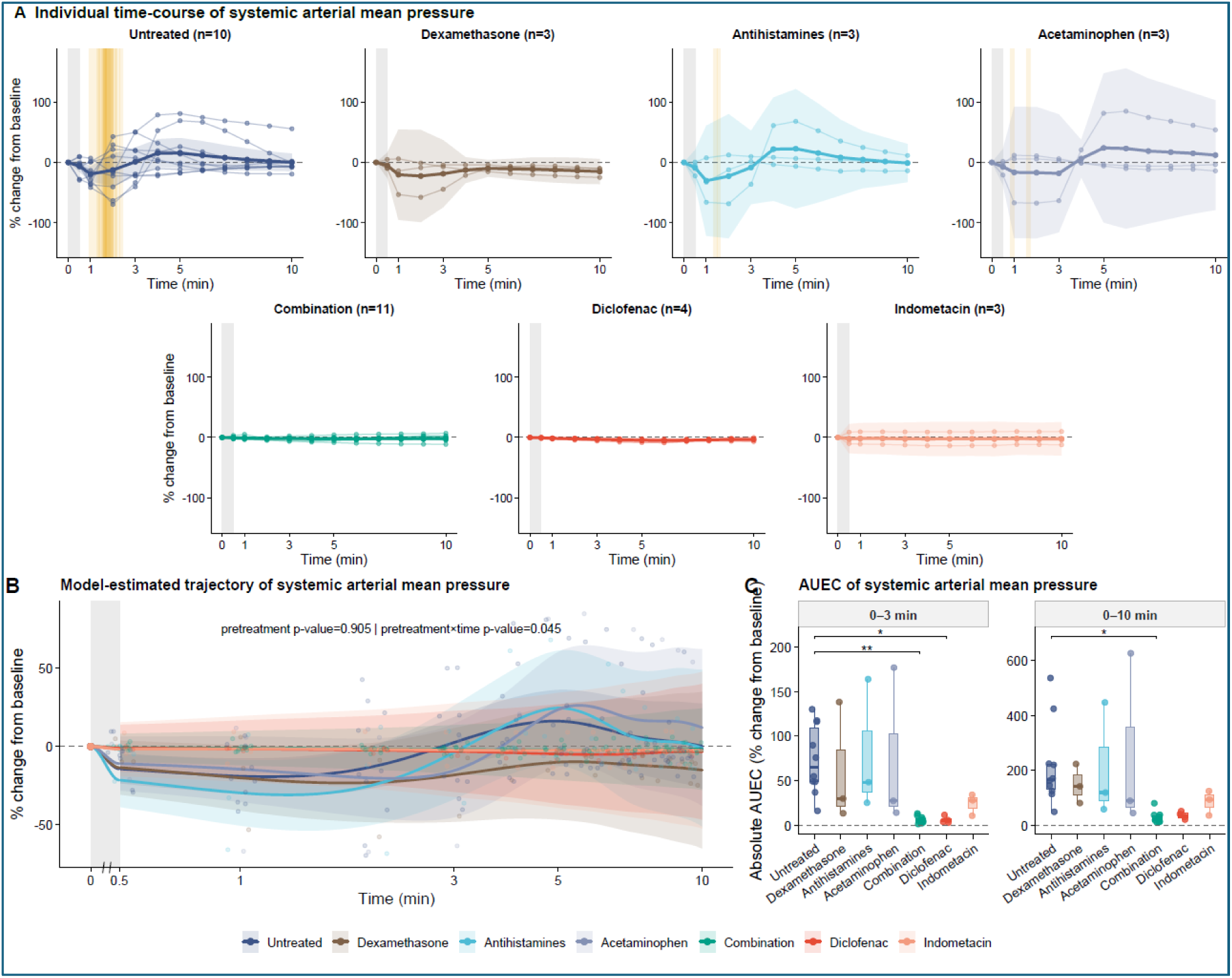
Effects of premedication regimens on Comirnaty-induced changes in mean SAP pressure expressed as percentage change from each animal’s baseline value. All other details are the same as in Fig. 3.

The wide and largely overlapping confidence bands in untreated animals and in those premedicated with dexamethasone, antihistamines, or acetaminophen indicate that these interventions did not substantially prevent SAP instability. By contrast, and in line with their effects on PAP, the triple combination of H1/H2 receptor blockade with diclofenac, as well as COX inhibitors alone, markedly suppressed the SAP alterations. This inhibitory effect was evident in the AUEC analysis during the early 0-3 min interval and remained apparent over the full 0-10 min observation period. Overall, these findings suggest that Comirnaty-induced systemic arterial pressure changes are closely linked to COX-dependent, most likely thromboxane-mediated, pulmonary vascular reactions rather than to histamine-dominated mechanisms alone.

### Effect of COX inhibitors on Comirnaty-induced pulse pressure changes

Because mSAP was partly constrained by the need for rescue treatment, we next reanalyzed the systemic arterial pressure response using pulse pressure, calculated as the difference between systolic and diastolic arterial pressure. Unlike the apparently variable mSAP response, pSAP revealed a marked, highly consistent fall after vaccine injection in control pigs and in animals premedicated with dexamethasone, antihistamines, or acetaminophen (Fig. 5A). This parameter integrates changes in stroke volume and systemic vascular resistance, and is therefore sensitive to both pulmonary vascular loading conditions and systemic vascular reactivity. It is also expected to change during severe vaccine-induced cardiovascular reactions and during catecholamine-supported stabilization, making it a more informative marker of the overall hemodynamic disturbance than mSAP alone.

**Figure 5.**
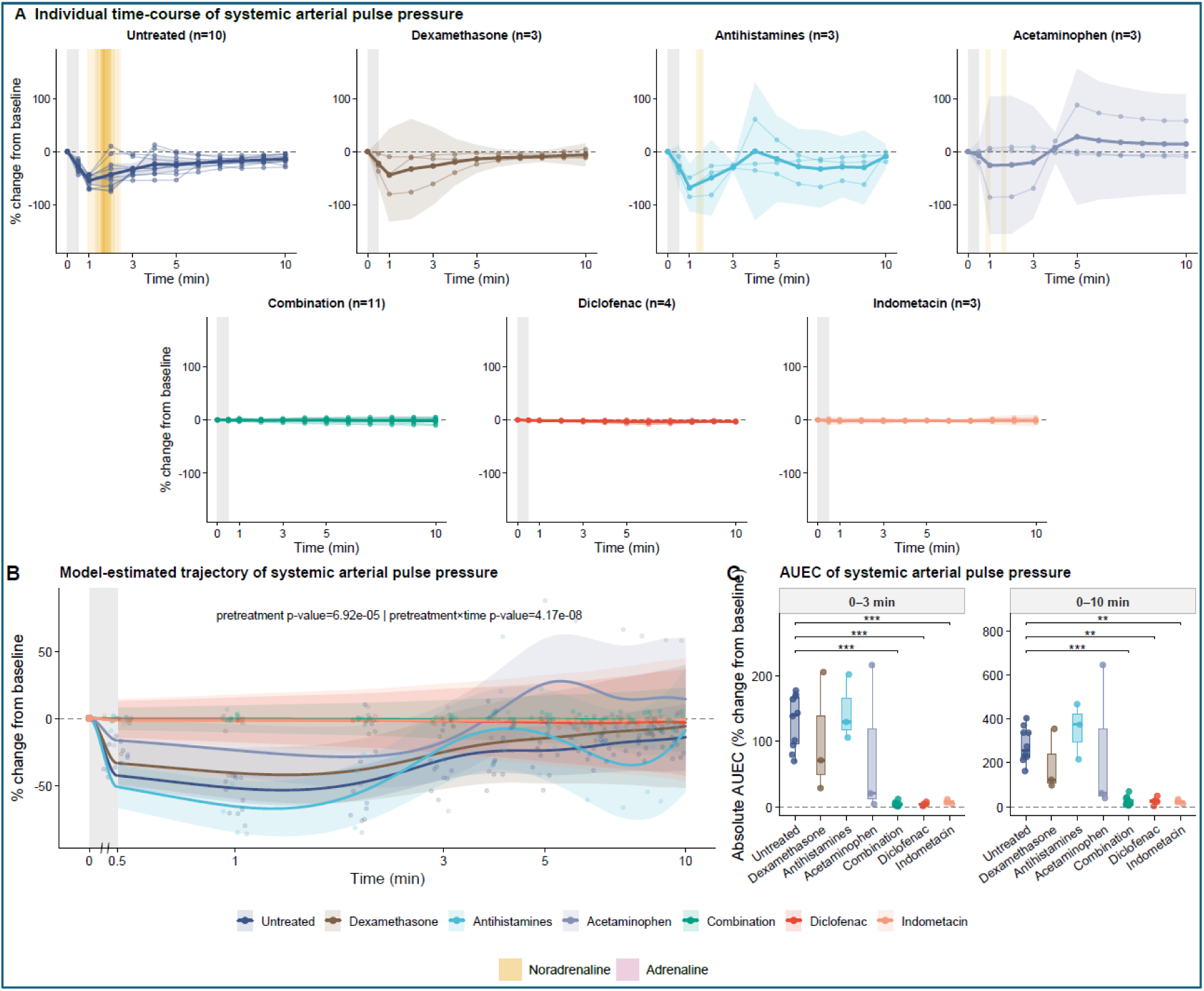
Effects of premedication protocols on Comirnaty-induced systemic arterial pulse pressure changes in anti-PEG hyperimmune pigs. The same experiment as shown in Figs. 3 and 4, except that the SAP changes were expressed as PP. All other conditions are the same as in Figs. 3 and 4.

Reanalyzing the data shown in Fig. 4 using pSAP instead of mSAP as an indicator of Comirnaty-induced systemic arterial pressure changes, Fig. 5A shows that the pSAP response was practically eliminated by COX inhibitors. Figures 5B and 6C further support this observation, showing the corresponding 95% confidence bands and AUC data, respectively. The marked reduction of pSAP changes by diclofenac and indomethacin parallels their effects on PAP (Fig. 3A-C), suggesting that inhibition of the pSAP response (Fig. 5A, C) and reduction in individual variability (Fig. 5B) are primarily due to suppression of pulmonary hypertension by COX inhibitors.

**Figure 6.**
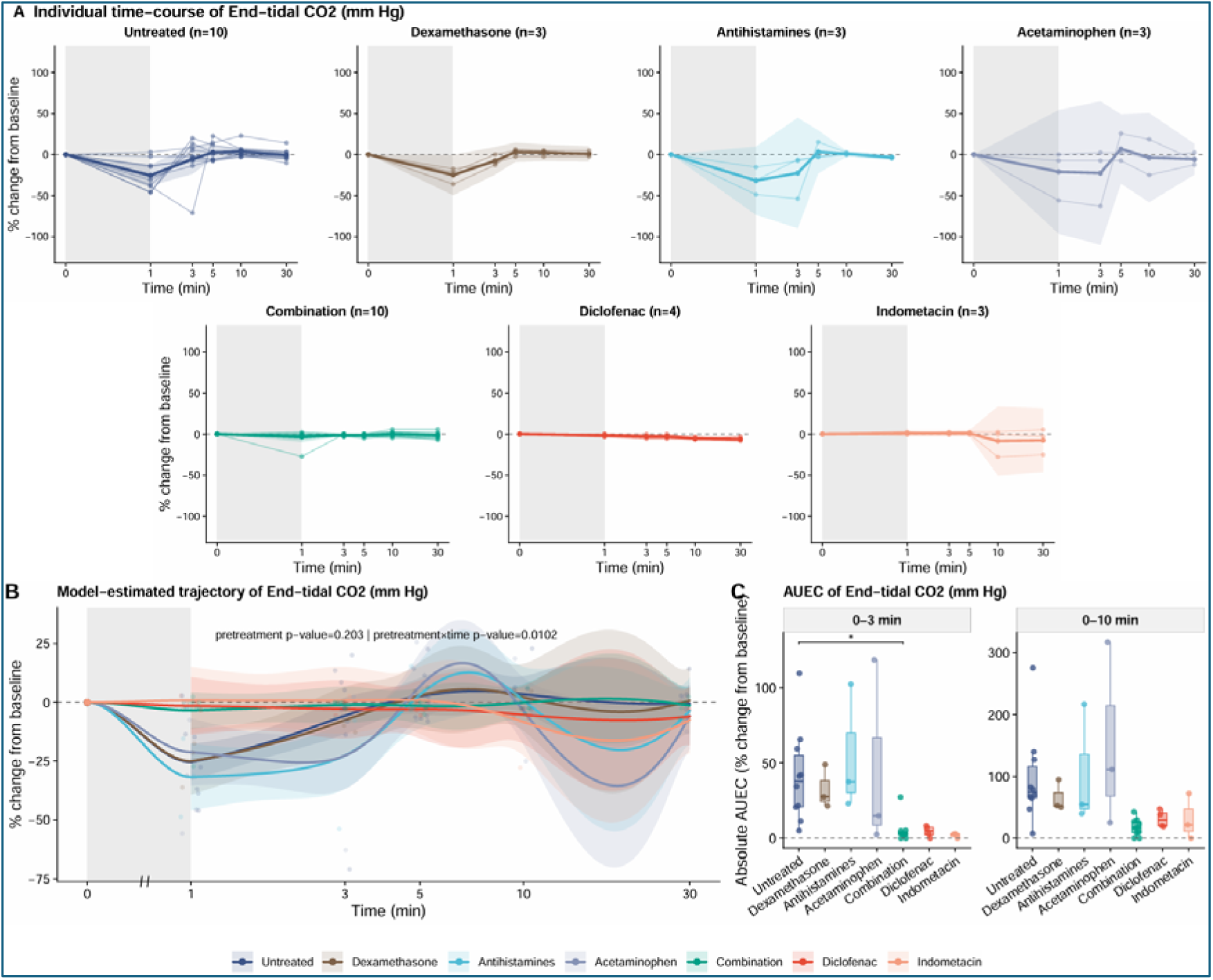
End-tidal partial pressure of CO□ (ETCO□) changes measured in the exhaled breath following Comirnaty administration. Datav from the same experiment as shown in Figs 3-5.

### Effect of COX inhibitors on Comirnaty-induced cardiopulmonary dysfunction

Figure 6 shows another effect of antihistamines and COX inhibitors, on end-tidal partial pressure of CO (ETCO□) measured in the exhaled breath at 1 min intervals following Comirnaty administration. It is a rapid, sensitive, noninvasive marker of cardiopulmonary function, providing an integrated readout of ventilation, cardiac output, and metabolism. As seen in the figure, the ETCO markedly decreased in the control (non-premedicated) and antihistamine group, indicating stagnation of gas exchange in the lung due to apnea and/or dyspnea. This decrease was completely prevented by both diclofenac and indomethacin. These findings further strengthen the conclusion that COX inhibition significantly attenuates the vaccine-induced cardiopulmonary dysfunction by inhibiting pulmonary vasoactivity.

### Clinical evidence for the efficacy enhancing effect of COX inhibitors on conventional premedication against liposome-induced infusion reactions

Pegylated nanomedicines in the form of liposomally-entrapped cytotoxic agents have been used in cancer therapy for nearly 30 years since the initial FDA approval of Doxil^®^]. Approximately10 years ago, we began including oral COX inhibitors (indomethacin or ibuprofen) along with antihistamines and dexamethasone in the premedication protocol applied at the Shaare Zedek Medical Center in Jerusalem, Israel. Since then, 362 patients (24 men and 338 women) have received Doxil^®^, 64 patients (24 men and 40 women) have received Promitil^®^, and only 3 patients have received Onivyde^®^ either as standard of care, as part of a clinical study or as compassionate treatment for cancer. Table 2 shows the distribution of these patients by sex, underlying disease and the incidence and severity of HSR/anaphylaxis rates caused by Doxil and Promitil infusion based on a retrospective assessment. Prior to the introduction of COX inhibitors, the literature suggests an approximate 7-11% incidence of HSR of variable severity for Doxil® [66].

**Table 2.**
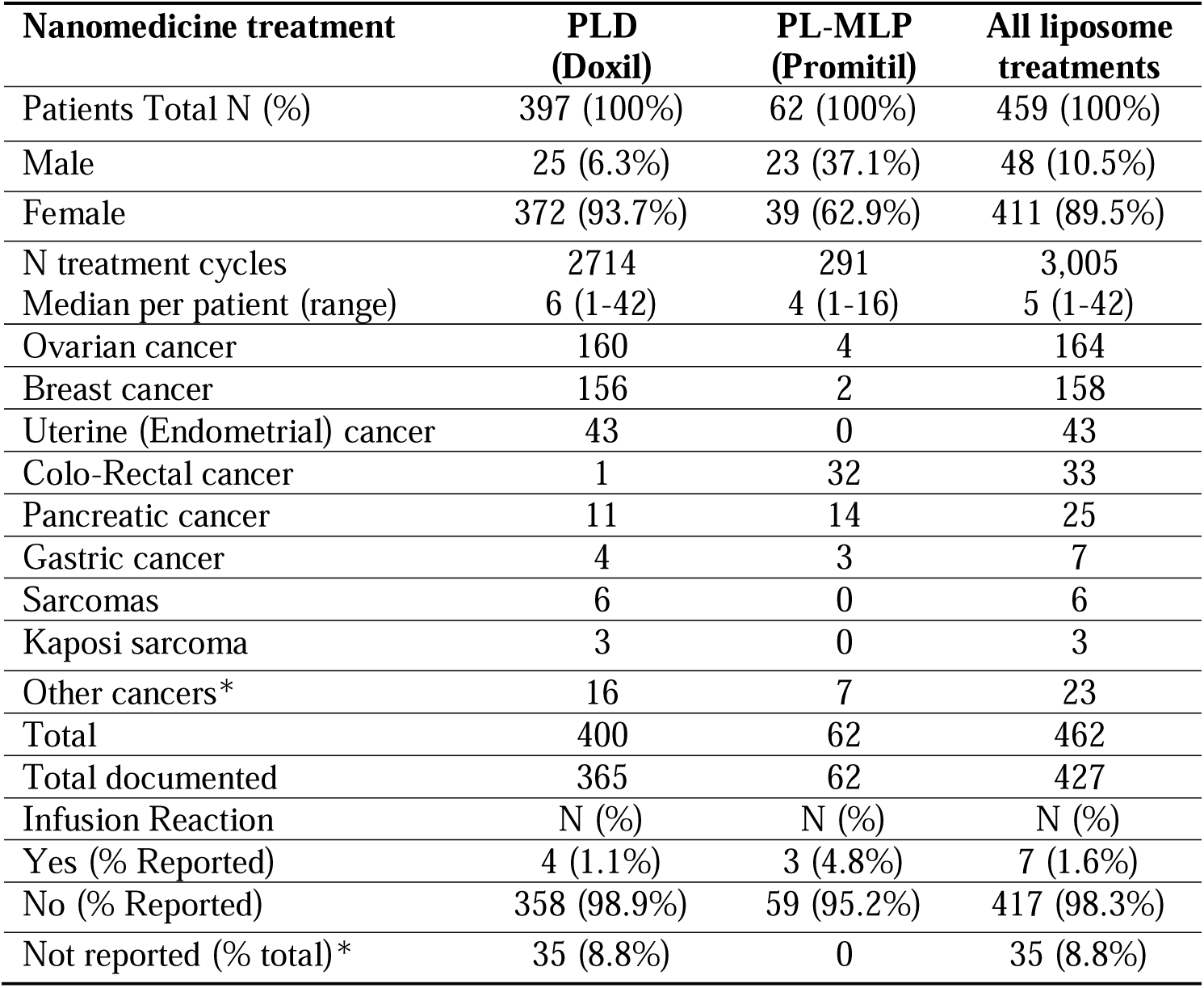
Distribution of patients treated with nanomedicines at Shaare Zede MC by sex and underlying disease.

| Nanomedicine treatment | PLD<br>(Doxil) | PL-MLP<br>(Promitil) | All liposome<br>treatments |
| --- | --- | --- | --- |
| Patients Total N (%) | 397 (100%) | 62 (100%) | 459 (100%) |
| Male | 25 (6.3%) | 23 (37.1%) | 48 (10.5%) |
| Female | 372 (93.7%) | 39 (62.9%) | 411 (89.5%) |
| N treatment cycles | 2714 | 291 | 3,005 |
| Median per patient (range) | 6 (1-42) | 4 (1-16) | 5 (1-42) |
| Ovarian cancer | 160 | 4 | 164 |
| Breast cancer | 156 | 2 | 158 |
| Uterine (Endometrial) cancer | 43 | 0 | 43 |
| Colo-Rectal cancer | 1 | 32 | 33 |
| Pancreatic cancer | 11 | 14 | 25 |
| Gastric cancer | 4 | 3 | 7 |
| Sarcomas | 6 | 0 | 6 |
| Kaposi sarcoma | 3 | 0 | 3 |
| Other cancers* | 16 | 7 | 23 |
| Total | 400 | 62 | 462 |
| Total documented | 365 | 62 | 427 |
| Infusion Reaction | N (%) | N (%) | N (%) |
| Yes (% Reported) | 4 (1.1%) | 3 (4.8%) | 7 (1.6%) |
| No (% Reported) | 358 (98.9%) | 59 (95.2%) | 417 (98.3%) |
| Not reported (% total)* | 35 (8.8%) | 0 | 35 (8.8%) |

Based on clinical records between 2015 and 2025. Symptomatic infusion reactions during Doxil/Promitil treatments were observed only in 7 cases out of a total of 427 patients documented (**1.6%**). The clinical manifestations were dyspnoea, back pain, facial/neck flushing, itching and rash. In 6 cases, the symptoms were Grade 2 (CTCAE Grades for Infusion-Related Reactions) and resolved with administration of IV hydrocortisone 100 mg and paracetamol PO 1 gr. After resuming treatment with slow infusion rate, patients completed the infusion successfully, except for one case in whom the reaction was severe (Grade 3) and it was decided not to resume the infusion and discontinue Doxil. There were only 3 cases treated with Onivyde (not shown), a low PEG-containing nanomedicine, with no infusion reactions reported.

*In a small group of Doxil-treated patients (8.8%), we have no information on patient tolerance of treatment. We refer to these patients as “Not reported”.

## Discussion

### Unmet medical need for reaction prevention of mRNA vaccine-induced anaphylaxis: Relevance beyond the pandemic

Hypersensitivity reactions, including anaphylaxis, are clinically relevant adverse events associated with mRNA-LNP vaccines. Indeed, in Pfizer’s 3-months cumulative post-authorization adverse event report anaphylaxis was identified as the only important “risks” of vaccination with Comirnaty” [44]. Although currently overshadowed by concerns over cardiac complications, anaphylaxis remains a primary safety issue, as reflected by its continued inclusion in the vaccine’s boxed warning label [1]. Anaphylaxis following mRNA vaccination is considered extremely rare; however, this designation applies only when judged by the standards of routinely prescribed drugs in general pharmacotherapy, particularly anticancer agents.

According to the Vaccine Adverse Event Reporting System (VAERS) in the USA [45], the combined incidence of anaphylaxis (per million doses) during the height of the vaccination campaign (December 2020 to May 2023) was approximately 60-fold higher than that observed for 12 influenza vaccines [46] over the same time. This disparity raises the question of why premedication against anaphylaxis has not become a standard procedure prior to vaccination in high-risk, atopic individuals with a history of severe allergic disease, whose proneness for vaccine induced anaphylaxis is much greater than that in the non-allergic population [8]. Currently, “only” those people are excluded from vaccination who has documented history of anaphylaxis to a previous dose of an mRNA COVID-19 vaccine, or any of its components [47]. Among the factors explaining this public health policy is the decline of vaccination campaign in any way, with vaccination in the USA now largely restricted to selected risk groups, thereby substantially reducing the absolute number of exposed individuals. Yet another contributing factor is the general oversight that mRNA-LNP vaccine formulations possess liposome-like immunological properties, resulting in reactions that share the same underlying mechanisms as liposomal infusion reactions [46,47]. Notably, LNPs do not contain a central aqueous compartment enclosed by phospholipid bilayers, as classical liposomes do; instead, they consist of a core in which mRNA is complexed with ionizable aminolipids [48–52]. Nevertheless, their exterior architecture, featuring a PEGylated phospholipid shell, confers liposome-like immunological behavior to LNPs.

### Utility of the anti-PEG antibody sensitized porcine model for the testing of premedication regimens against vaccine-induced anaphylaxis

Anaphylaxis, affecting both animals and humans, cannot be adequately studied using in vitro, ex vivo, or in silico approaches. Among animal models, rodents and other commonly used laboratory species are typically insensitive to many human-reactogenic drugs and therefore provide limited predictive value for HSR risk in humans [53, 54]. In contrast, treatment of pigs, sheep, and other large animals can reproduce allergy-like syndromes that closely resemble those observed in hypersensitive humans. Among these, the porcine CARPA model [55], particularly its anti-PEG antibody-sensitized variant [14], has emerged as the most sensitive and reproducible approach for reactogenicity assessment, including vaccine-induced reactions.

### The mechanism of vaccine reactions

Vaccine-induced HSRs closely mirror those triggered by liposomal nanomedicines in their clinical, hemodynamic, and immunological features, consistent with CARPA as a shared mechanism. In both settings, rapid C activation, frequently mediated by anti-PEG antibodies, induces acute cardiopulmonary distress through the release of vasoactive mediators, reflecting an unusual stress reaction via the immune-cardiovascular system axis [11]. Experimental evidence identifying C and secondary innate immune cell activation as the primary drivers of these reactions [7, 14, 55] provides a unifying explanation for HSRs to both mRNA-LNP vaccines and PEGylated liposomal drugs.

### The effects of antihistamines

Levocetirizine is used as a second-generation selective peripheral H1 histamine receptor antagonist blocking histamine-mediated allergic reactions, while famotidine, a guanidylthiazole derivative that inhibits H2 histamine receptors, is commonly used in human medicine. It inhibits cytokine release by blocking H2 histamine receptors on immune system cells (neutrophil granulocytes, T cells, mast cells, etc.).

Although not critical from a life-threatening perspective, antihistamines did exert differential effects on systemic and pulmonary blood pressures. Specifically, they increased the variability of SAP, while reducing the variability of PAP. This is consistent with the fact that SAP is determined by both pulmonary blood flow and peripheral vascular resistance, whereas PAP mainly reflects pulmonary vasotonicity. Accordingly, the pronounced inter-individual variability in mean SAP likely reflects differing contributions of pulmonary vascular obstruction and peripheral vasomotor responses to antihistamines. Accordingly, pulmonary hypertension-induced systemic hypotension may be superimposed on peripheral vascular effects, resulting in wide SAP variability across animals. In essence, Comirnaty exerts differential effects on the pulmonary and systemic vascular compartments. When peripheral vascular influences are separated from pulmonary obstruction, inhibition of pulmonary hypertension should also be detectable in the initial SAP decline, best captured by pulse pressure. Consistent with this interpretation, reanalysis using pulse pressure as the SAP endpoint (Figure 6) showed that COX inhibitors significantly inhibited pulse pressure changes to a degree comparable to its effect on pulmonary hypertension.

### History of premedication against liposome reactions

Premedication against liposome-induced HSRs was introduced following early clinical experience with i.v. administered radiocontrast media [56] and drugs, which revealed frequent acute infusion reactions [57–60]. These empirically designed, so-called “kitchen-sink” regimens were largely exploratory and not based on systematic mechanistic analysis. Over time, guided primarily by accumulated clinical experience, they evolved to include the ingredients and applications listed in Table 3.

**Table 3.**
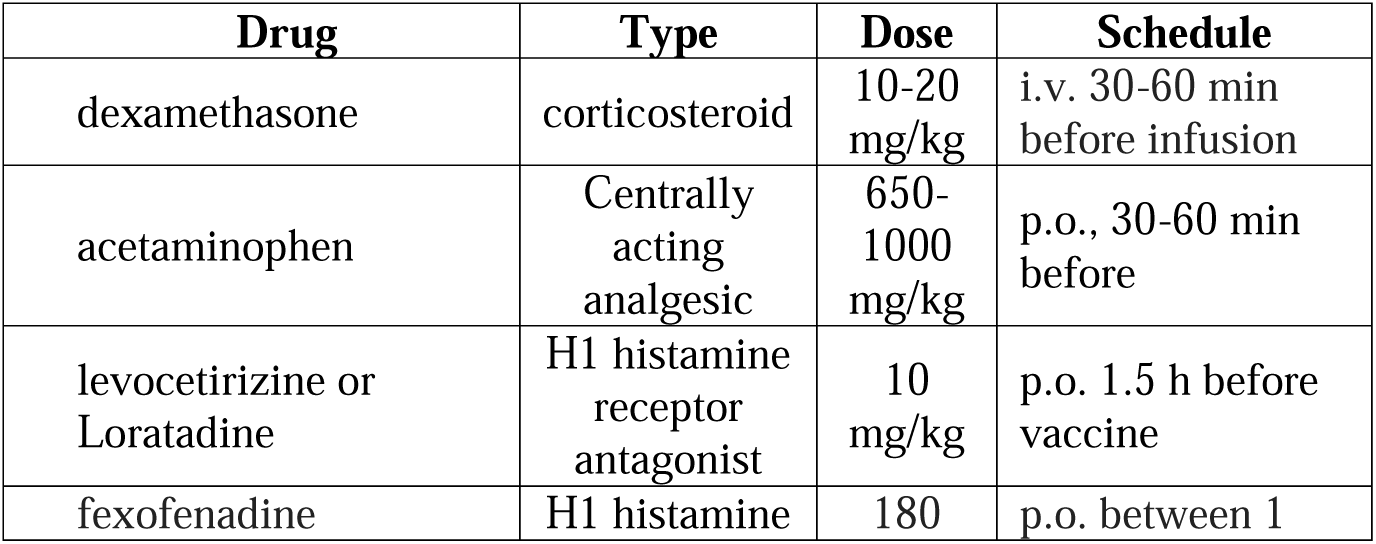
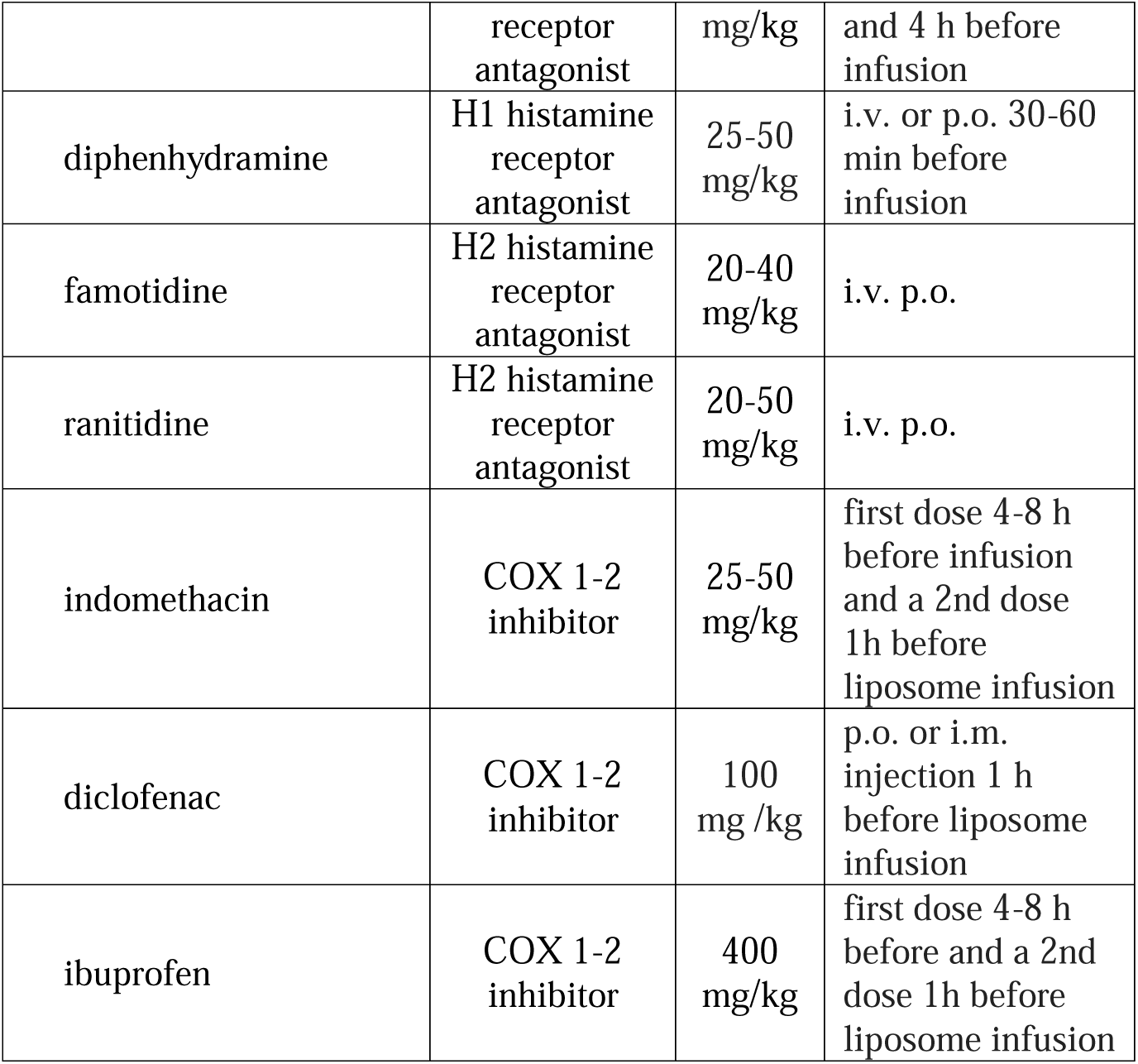
Clinically and experimentally used premedication drugs and their dosing regimens.

It is likely that COX inhibitors, a well-studied class of anti-inflammatory drugs, have been largely excluded from most premedication protocols because inflammation is not perceived as a hallmark of HSR’s.

### COX inhibition emerges as the most effective premedication strategy

Indomethacin and diclofenac are classical nonsteroidal anti-inflammatory drugs (NSAID) that directly bind to and inhibits the COX active site, thereby blocking the conversion of arachidonic acid into prostaglandins and thromboxanes (PGE□, PGI□, TXA□). Unlike classical COX inhibitors, acetaminophen also inhibits COX activity and has anti-inflammatory and antipyretic effects; however, its action is exerted predominantly within the central nervous system as a reductive COX inhibitor, interacting with the peroxidase site, not the cyclooxygenase site of the enzyme [61]. Nevertheless, our data with acetaminophen did suggest a more moderate suppressive effect on PAP and SAP changes (Fig. 5,6), at least.

Cyclooxygenase inhibitors were sporadically included in early premedication regimens for infusion reactions, particularly in oncology settings, but they were not a standard or uniformly applied component of liposome-premedication protocols. Consequently, unlike corticosteroids and antihistamines, COX inhibitors were never rigorously or independently validated as preventive agents for liposome-induced HSRs. A milestone in this regard was a porcine study published in 1999 [62], in which the CARPA mechanism of anaphylactoid reactions to liposomes was first described. In that work, indomethacin was shown to completely prevent the characteristic rise of PAP [62].

Since then, to our knowledge, the only clinical oncology center that has consistently incorporated and reported [16] on the specific efficacy of COX inhibitors as part of premedication protocols is Shaare Zedek Medical Center, through the work of a co-author of this study (AG).

### Broader implications of COX inhibitors preventing anaphylaxis

Altogether, optimization of premedication regimens may enable safer and broader use not only of mRNA vaccines but also of all liposome- and LNP-containing reactogenic nanomedicines. The anti-PEG antibody-sensitized model increases susceptibility to anaphylactic reactions induced by PEGylated nanoparticles, thereby providing a robust analytical system for elucidating the mechanisms of this immune reactivity and for exploring effective preventive strategies.

Although dexamethasone, H1 and H2 antihistamines, and the centrally acting analgesic acetaminophen modulated vaccine-induced reactions, only COX inhibitors proved effective in fully eliminating all reaction symptoms. This observation raises the question of why, and under what circumstances, non-COX inhibitors should be included in premedication regimens. A particular concern is the immunosuppressive effect of dexamethasone, which may counteract the efficacy of vaccines or other immune stimulatory therapies. If confirmed for other reactogenic drugs and clinical protocols, these findings suggest that premedication regimens could be simplified to COX-inhibitor monotherapy, potentially reducing costs, treatment burden, and patient concern.

## Acknowledgment

The technical assistance of Henriett Biro, Gabriella Juhász, Benjamin Prokaj, Ádám Steiner, Zsombor Fáskerti, and Zoltán Manhertz is gratefully acknowledged.

## Supplemental Information

**Figure S1.**
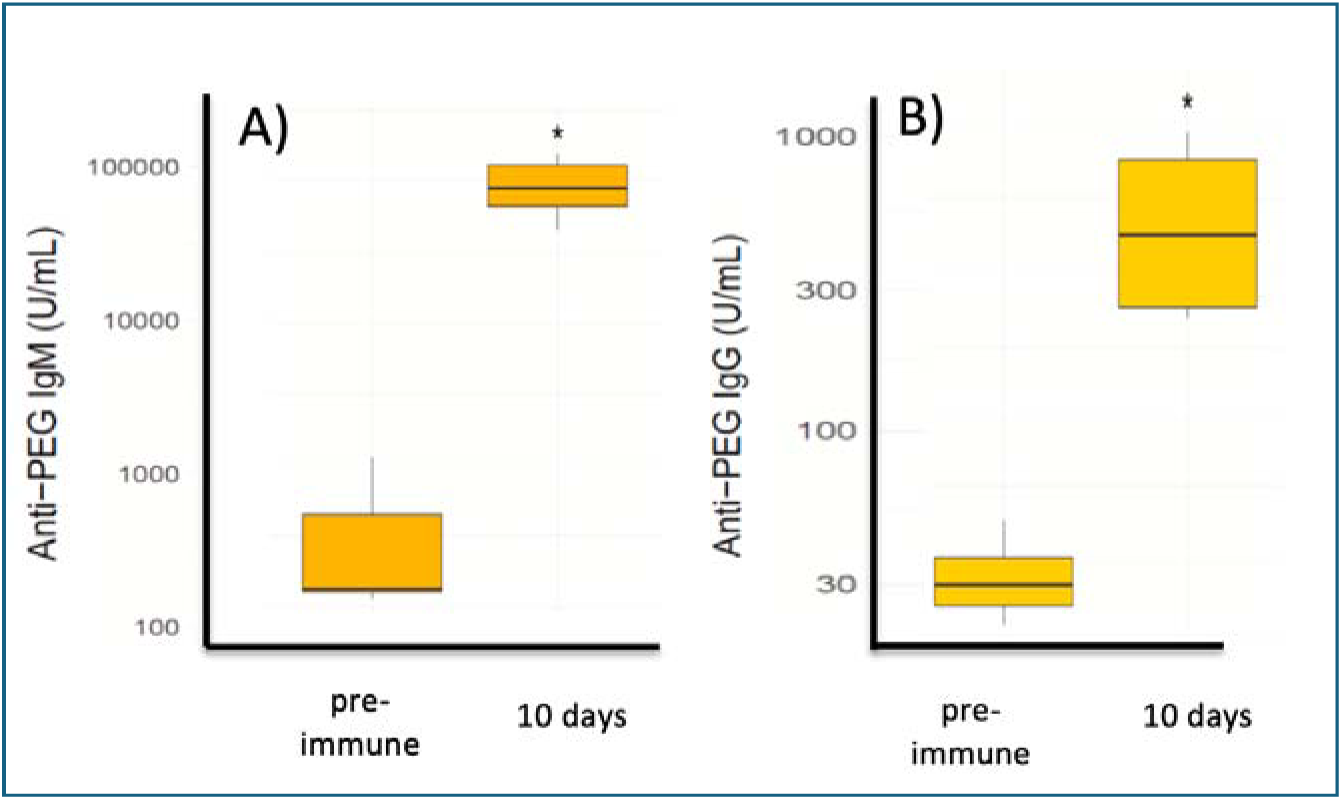
Rise of anti-PEG IgM and IgG in pig blood after injection of Doxebo. Plasma anti-PEG antibody levels were measured before immunization (“pre-immune”) and 9–10 days after treatment with Doxebo. Box plots show anti-PEG IgM (A) and anti-PEG IgG (B) concentrations expressed as U/mL on a logarithmic scale. Boxes indicate the interquartile range, horizontal lines show the median, and whiskers indicate the data range. The figure has been reproduced from [14] with permission. *p < 0.05 versus pre-immune.

## References

1. FDA, US. “Prescribing Information: Comirnaty® (Covid-19 Vaccine, Mrna) Injectable Suspension, for Intramuscular Use “ https://www.fda.gov/media/151707/download?utm_source=chatgpt.com (2025).

2. Miller, E. R., M. M. McNeil, P. L. Moro, J. Duffy, and J. R. Su. “The Reporting Sensitivity of the Vaccine Adverse Event Reporting System (Vaers) for Anaphylaxis and for Guillain-Barre Syndrome.” Vaccine 38, no. 47 (2020): 7458–63.

3. Shimabukuro, T. T., M. Cole, and J. R. Su. “Reports of Anaphylaxis after Receipt of Mrna Covid-19 Vaccines in the Us-December 14, 2020-January 18, 2021.” JAMA 325, no. 11 (2021): 1101–02.

4. Jaggers, J., and A. R. Wolfson. “Mrna Covid-19 Vaccine Anaphylaxis: Epidemiology, Risk Factors, and Evaluation.” Curr Allergy Asthma Rep 23, no. 3 (2023): 195–200.

5. Khalid, M. B., and P. A. Frischmeyer-Guerrerio. “The Conundrum of Covid-19 Mrna Vaccine-Induced Anaphylaxis.” J Allergy Clin Immunol Glob 2, no. 1 (2023): 1–13.

6. Song, J., D. Su, H. Wu, and J. Guo. “Implications of Anaphylaxis Following Mrna-Lnp Vaccines: It Is Urgent to Eliminate Peg and Find Alternatives.” Pharmaceutics 17, no. 6 (2025).

7. Kozma, G. T., T. Meszaros, P. Berenyi, R. Facsko, Z. Patko, C. Z. Olah, A. Nagy, T. G. Fulop, K. A. Glatter, T. Radovits, B. Merkely, and J. Szebeni. “Role of Anti-Polyethylene Glycol (Peg) Antibodies in the Allergic Reactions to Peg-Containing Covid-19 Vaccines: Evidence for Immunogenicity of Peg.” Vaccine 41, no. 31 (2023): 4561–70.

8. Shavit, R., R. Maoz-Segal, M. Iancovici-Kidon, I. Offengenden, S. Haj Yahia, D. Machnes Maayan, Y. Lifshitz-Tunitsky, S. Niznik, S. Frizinsky, M. Deutch, E. Elbaz, H. Genaim, G. Rahav, I. Levy, A. Belkin, G. Regev-Yochay, A. Afek, and N. Agmon-Levin. “Prevalence of Allergic Reactions after Pfizer-Biontech Covid-19 Vaccination among Adults with High Allergy Risk.” JAMA Netw Open 4, no. 8 (2021): e2122255.

9. Warren, C. M., T. T. Snow, A. S. Lee, M. M. Shah, A. Heider, A. Blomkalns, B. Betts, A. S. Buzzanco, J. Gonzalez, R. S. Chinthrajah, E. Do, I. Chang, D. Dunham, G. Lee, R. O’Hara, H. Park, M. H. Shamji, L. Schilling, S. B. Sindher, D. Sisodiya, E. Smith, M. Tsai, S. J. Galli, C. Akdis, and K. C. Nadeau. “Assessment of Allergic and Anaphylactic Reactions to Mrna Covid-19 Vaccines with Confirmatory Testing in a Us Regional Health System.” JAMA Netw Open 4, no. 9 (2021): e2125524.

10. Szebeni, J. “Complement Activation-Related Pseudoallergy: A New Class of Drug-Induced Acute Immune Toxicity.” Toxicology 216, no. 2-3 (2005): 106–21.

11. Szebeni, J. “Complement Activation-Related Pseudoallergy: A Stress Reaction in Blood Triggered by Nanomedicines and Biologicals.” Molecular immunology 61, no. 2 (2014): 163–73.

12. Saxena, S., S. Sharma, G. Kumar, and S. Thakur. “Unravelling the Complexity of Carpa: A Review of Emerging Advancements in Therapeutic Strategies.” Arch Dermatol Res 317, no. 1 (2025): 439.

13. Dezsi, L., T. Meszaros, G. Kozma, H. Velkei M, C. Z. Olah, M. Szabo, Z. Patko, T. Fulop, M. Hennies, M. Szebeni, B. A. Barta, B. Merkely, T. Radovits, and J. Szebeni. “A Naturally Hypersensitive Porcine Model May Help Understand the Mechanism of Covid-19 Mrna Vaccine-Induced Rare (Pseudo) Allergic Reactions: Complement Activation as a Possible Contributing Factor.” Geroscience 44, no. 2 (2022): 597–618.

14. Barta, B. A., T. Radovits, A. B. Dobos, G. Tibor Kozma, T. Meszaros, P. Berenyi, R. Facsko, T. Fulop, B. Merkely, and J. Szebeni. “Comirnaty-Induced Cardiopulmonary Distress and Other Symptoms of Complement-Mediated Pseudo-Anaphylaxis in a Hyperimmune Pig Model: Causal Role of Anti-PEG Antibodies.” Vaccine X 19 (2024): 100497.

15. Szebeni, J., G. Storm, J. Y. Ljubimova, M. Castells, E. J. Phillips, K. Turjeman, Y. Barenholz, D. J. A. Crommelin, and M. A. Dobrovolskaia. “Applying Lessons Learned from Nanomedicines to Understand Rare Hypersensitivity Reactions to Mrna-Based Sars-Cov-2 Vaccines.” Nat Nanotechnol 17, no. 4 (2022): 337–46.

16. Gabizon, A., and J. Szebeni. “Complement Activation: A Potential Threat on the Safety of Poly(Ethylene Glycol)-Coated Nanomedicines.” ACS Nano 14, no. 7 (2020): 7682–88.

17. Chanan-Khan, A., J. Szebeni, S. Savay, L. Liebes, N.M. Rafique, C.R. Alving, and F.M. Muggia. “Complement Activation Following First Exposure to Pegylated Liposomal Doxorubicin (Doxil): Possible Role in Hypersensitivity Reactions.” Ann Oncol 14 (2003): 1430–37.

18. Szebeni, J., P. Bedocs, R. Urbanics, R. Bunger, L. Rosivall, M. Toth, and Y. Barenholz. “Prevention of Infusion Reactions to Pegylated Liposomal Doxorubicin Via Tachyphylaxis Induction by Placebo Vesicles: A Porcine Model.” J Control Release 160, no. 2 (2012): 382–7.

19. Bavli, Y., I. Winkler, B. M. Chen, S. Roffler, R. Cohen, J. Szebeni, and Y. Barenholz. “Doxebo (Doxorubicin-Free Doxil-Like Liposomes) Is Safe to Use as a Pre-Treatment to Prevent Infusion Reactions to Pegylated Nanodrugs.” J Control Release 306 (2019): 138–48.

20. Fulop, T., G. T. Kozma, I. Vashegyi, T. Meszaros, L. Rosivall, R. Urbanics, G. Storm, J. M. Metselaar, and J. Szebeni. “Liposome-Induced Hypersensitivity Reactions: Risk Reduction by Design of Safe Infusion Protocols in Pigs.” J Control Release 309 (2019): 333–38.

21. Walsh, T.J., J.L. Goodman, P. Papas, I. Bekersky, D.N. Buell, M. Roden, J. Barrett, and E.J. Anaissie. “Safety, Tolerance, and Pharmacokinetics of High-Dose Liposomal Amphotericin B (Ambisome) in Patients Infected with Aspergillus Species and Other Filamentous Fungi: Maximum Tolerated Dose Study.” Antimicrobial Agents and Chemotherapy 45 (2001): 3487–96.

22. Roden, M. M., L. D. Nelson, T. A. Knudsen, P. F. Jarosinski, J. M. Starling, S. E. Shiflett, K. Calis, R. DeChristoforo, G. R. Donowitz, D. Buell, and T. J. Walsh. “Triad of Acute Infusion-Related Reactions Associated with Liposomal Amphotericin B: Analysis of Clinical and Epidemiological Characteristics.” Clin Infect Dis 36, no. 10 (2003): 1213–20.

23. Ringden, O., E. Andstrom, M. Remberger, B. M. Svahn, and J. Tollemar. “Allergic Reactions and Other Rare Side-Effects of Liposomal Amphotericin.” Lancet 344, no. 8930 (1994): 1156–7.

24. Tahover, E., R. Bar-Shalom, E. Sapir, R. Pfeffer, I. Nemirovsky, Y. Turner, M. Gips, P. Ohana, B. W. Corn, A. Z. Wang, and A. A. Gabizon. “Chemo-Radiotherapy of Oligometastases of Colorectal Cancer with Pegylated Liposomal Mitomycin-C Prodrug (Promitil): Mechanistic Basis and Preliminary Clinical Experience.” Front Oncol 8 (2018): 544.

25. Gabizon, A., H. Shmeeda, E. Tahover, G. Kornev, Y. Patil, Y. Amitay, P. Ohana, E. Sapir, and S. Zalipsky. “Development of Promitil(R), a Lipidic Prodrug of Mitomycin C in Pegylated Liposomes: From Bench to Bedside.” Adv Drug Deliv Rev 154-155 (2020): 13–26.

26. Jaiswal, V., K. Kalra, N. Deb, and J. Mattumpuram. “Cardiovascular Safety of Patisiran among Transthyretin Cardiac Amyloidosis: A Meta-Analysis.” Am J Cardiovasc Drugs 25, no. 1 (2025): 125–27.

27. Huang, X., C. Sun, H. Chen, C. Zhao, and J. Lin. “Efficacy and Safety of Patisiran for Attrv-Pn: A Systematic Review and Meta-Analysis.” Ther Adv Neurol Disord 17 (2024): 17562864241273079.

28. Karimi, M. A., F. Esmaeilpour Moallem, M. S. Gholami Chahkand, E. Azarm, M. J. Emami Kazemabad, and P. A. Dadkhah. “Assessing the Effectiveness and Safety of Patisiran and Vutrisiran in Attrv Amyloidosis with Polyneuropathy: A Systematic Review.” Front Neurol 15 (2024): 1465747.

29. Hagemeister, F., M. A. Rodriguez, S. R. Deitcher, A. Younes, L. Fayad, A. Goy, N. H. Dang, A. Forman, P. McLaughlin, L. J. Medeiros, B. Pro, J. Romaguera, F. Samaniego, J. A. Silverman, A. Sarris, and F. Cabanillas. “Long Term Results of a Phase 2 Study of Vincristine Sulfate Liposome Injection (Marqibo((R) Substituted for Non-Liposomal Vincristine in Cyclophosphamide, Doxorubicin, Vincristine, Prednisone with or without Rituximab for Patients with Untreated Aggressive Non-Hodgkin Lymphomas.” Br J Haematol 162, no. 5 (2013): 631–8.

30. Silverman, J. A., L. Reynolds, and S. R. Deitcher. “Pharmacokinetics and Pharmacodynamics of Vincristine Sulfate Liposome Injection (Vsli) in Adults with Acute Lymphoblastic Leukemia.” J Clin Pharmacol 53, no. 11 (2013): 1139–45.

31. Gao, Y., Y. Xie, L. Liu, H. Xue, M. Hou, M. Zhang, Z. Zhou, P. Guo, H. Yao, Z. Shao, X. Xie, and J. Zhu. “Efficacy and Safety of Cyclophosphamide, Doxorubicin, Vincristine, and Prednisone Regimen with Pegylated Liposomal Doxorubicin+/-Rituximab in Treating Diffuse Large B-Cell Lymphoma.” Minerva Med 112, no. 2 (2021): 310–12.

32. Wang-Gillam, A., C. P. Li, G. Bodoky, A. Dean, Y. S. Shan, G. Jameson, T. Macarulla, K. H. Lee, D. Cunningham, J. F. Blanc, R. A. Hubner, C. F. Chiu, G. Schwartsmann, J. T. Siveke, F. Braiteh, V. Moyo, B. Belanger, N. Dhindsa, E. Bayever, D. D. Von Hoff, L. T. Chen, and Napoli- Study Group. “Nanoliposomal Irinotecan with Fluorouracil and Folinic Acid in Metastatic Pancreatic Cancer after Previous Gemcitabine-Based Therapy (Napoli-1): A Global, Randomised, Open-Label, Phase 3 Trial.” Lancet 387, no. 10018 (2016): 545–57.

33. Wang-Gillam, A., D. Von Hoff, J. Siveke, R. Hubner, B. Belanger, J. M. Pipas, and L. T. Chen. “Nanoliposomal Irinotecan in the Clinical Practice Guideline for Metastatic Pancreatic Cancer: Applicability to Clinical Situations.” J Clin Oncol 35, no. 6 (2017): 689–90.

34. Carnevale, J., and A. H. Ko. “Mm-398 (Nanoliposomal Irinotecan): Emergence of a Novel Therapy for the Treatment of Advanced Pancreatic Cancer.” Future Oncol 12, no. 4 (2016): 453–64.

35. Chiang, N. J., J. Y. Chang, Y. S. Shan, and L. T. Chen. “Development of Nanoliposomal Irinotecan (Nal-Iri, Mm-398, Pep02) in the Management of Metastatic Pancreatic Cancer.” Expert Opin Pharmacother 17, no. 10 (2016): 1413–20.

36. Wang, W., Y. Yi, Y. Jia, X. Dong, J. Zhang, X. Song, and Y. Song. “Neoadjuvant Chemotherapy with Liposomal Paclitaxel Plus Platinum for Locally Advanced Esophageal Squamous Cell Cancer: Results from a Retrospective Study.” Thorac Cancer 13, no. 6 (2022): 824–31.

37. Li, R., H. Liang, J. Li, Z. Shao, D. Yang, J. Bao, K. Wang, W. Xi, Z. Gao, R. Guo, and X. Mu. “Paclitaxel Liposome (Lipusu) Based Chemotherapy Combined with Immunotherapy for Advanced Non-Small Cell Lung Cancer: A Multicenter, Retrospective Real-World Study.” BMC Cancer 24, no. 1 (2024): 107.

38. Mohammad-Jafari, K., S. M. Naghib, and M. R. Mozafari. “Liposomal Nanoformulation-Encapsulated Paclitaxel for Reducing Chemotherapy Side Effects in Lung Cancer Treatments: Recent Advances and Future Outlooks.” Curr Med Chem 32, no. 25 (2025): 5155–79.

39. Stathopoulos, G. P., T. Boulikas, M. Vougiouka, G. Deliconstantinos, S. Rigatos, E. Darli, V. Viliotou, and J. G. Stathopoulos. “Pharmacokinetics and Adverse Reactions of a New Liposomal Cisplatin (Lipoplatin): Phase I Study.” Oncol Rep 13, no. 4 (2005): 589–95.

40. Stathopoulos, G. P., and T. Boulikas. “Lipoplatin Formulation Review Article.” J Drug Deliv 2012 (2012): 581363.

41. Cook, G., and I. M. Franklin. “Adverse Drug Reactions Associated with the Administration of Amphotericin B Lipid Complex (Abelcet).” Bone Marrow Transplant 23, no. 12 (1999): 1325–6.

42. Milosevits, G., T. Meszaros, E. Orfi, T. Bakos, M. Garami, G. Kovacs, L. Dezsi, P. Hamar, B. Gyorffy, A. Szabo, G. Szenasi, and J. Szebeni. “Complement-Mediated Hypersensitivity Reactions to an Amphotericin B-Containing Lipid Complex (Abelcet) in Pediatric Patients and Anesthetized Rats: Benefits of Slow Infusion.” Nanomedicine 34 (2021): 102366.

43. Gallis, H. A., R. H. Drew, and W. W. Pickard. “Amphotericin B: 30 Years of Clinical Experience.” Rev Infect Dis 12, no. 2l 308–29 (1990)

44. Worldwide Safety. “Cumulative Analysis of Post-Authorization Adverse Event Reports of Pf-07302048 (Bnt162b2) Received through 28-Feb-2021.” https://phmpt.org/wp-content/uploads/2021/11/5.3.6-postmarketing-experience.pdf?fbclid=IwAR2tWI7DKw0cc2lj8 (2021).

45. OpenVAERS. “Vaers Covid Vaccine Adverse Event Reports.” https://openvaers.com/covid-data https://openvaers.com/covid-data (2024).

46. Szebeni, J. “Expanded Spectrum and Increased Incidence of Adverse Events Linked to Covid-19 Genetic Vaccines: New Concepts on Prophylactic Immuno-Gene Therapy, Iatrogenic Orphan Disease, and Platform-Inherent Challenges.” Pharmaceutics 17, no. 4 10.3390/pharmaceutics17040450 (2025)

47. Szebeni, J. “Unique Features and Collateral Immune Effects of mRNA-LNP COVID-19 Vaccines: Plausible Mechanisms of Adverse Events and Complications“ Pharmaceutics;17(10):1327. doi: 10.3390/pharmaceutics17101327 (2025)

48. Pfizer, and Biontech. “Comirnaty Original/Omicron Ba.4-5 Dispersion for Injection.” https://labeling.pfizer.com/ShowLabeling.aspx?id=19823 (2023).

49. Kulkarni, J. A., M. M. Darjuan, J. E. Mercer, S. Chen, R. van der Meel, J. L. Thewalt, Y. Y. C. Tam, and P. R. Cullis. “On the Formation and Morphology of Lipid Nanoparticles Containing Ionizable Cationic Lipids and Sirna.” ACS Nano 12, no. 5 (2018): 4787–95.

50. Sebastiani, F., M. Yanez Arteta, M. Lerche, L. Porcar, C. Lang, R. A. Bragg, C. S. Elmore, V. R. Krishnamurthy, R. A. Russell, T. Darwish, H. Pichler, S. Waldie, M. Moulin, M. Haertlein, V. T. Forsyth, L. Lindfors, and M. Cardenas. “Apolipoprotein E Binding Drives Structural and Compositional Rearrangement of Mrna-Containing Lipid Nanoparticles.” ACS Nano 15, no. 4 (2021): 6709–22.

51. Szebeni, J., B. Kiss, T. Bozo, K. Turjeman, Y. Levi-Kalisman, Y. Barenholz, and M. Kellermayer. “Insights into the Structure of Comirnaty Covid-19 Vaccine: A Theory on Soft, Partially Bilayer-Covered Nanoparticles with Hydrogen Bond-Stabilized Mrna-Lipid Complexes.” ACS Nano 17, no. 14 (2023): 13147–57.

52. Cheng, M. H. Y., J. Leung, Y. Zhang, C. Strong, G. Basha, A. Momeni, Y. Chen, E. Jan, A. Abdolahzadeh, X. Wang, J. A. Kulkarni, D. Witzigmann, and P. R. Cullis. “Induction of Bleb Structures in Lipid Nanoparticle Formulations of Mrna Leads to Improved Transfection Potency.” Adv Mater 35, no. 31 (2023): e2303370.

53. Szebeni, J., C. R. Alving, L. Rosivall, R. Bunger, L. Baranyi, P. Bedocs, M. Toth, and Y. Barenholz. “Animal Models of Complement-Mediated Hypersensitivity Reactions to Liposomes and Other Lipid-Based Nanoparticles.” J Liposome Res 17, no. 2 (2007): 107–17.

54. Szebeni, J. “Evaluation of the Acute Anaphylactoid Reactogenicity of Nanoparticle-Containing Medicines and Vaccines Using the Porcine Carpa Model.” Methods Mol Biol 2789 (2024): 229–43.

55. Dézsi, László, Gábor Kökény, Gábor Szénási, Csaba Révész, Tamás Mészáros, Bálint A Barta, Réka Facsko, Anna Szilasi, Tamás Bakos, and Gergely T Kozma. “Acute Anaphylactic and Multiorgan Inflammatory Effects of Comirnaty in Pigs: Evidence of Spike Protein mRNA Transfection and Paralleling Inflammatory Cytokine Upregulation.” BioRxiv, https://www.biorxiv.org/content/10.1101/2025.06.07.658379v2 2025.06. 07.658379, no. 658379 (2025).

56. Greenberger, P. A., R. Patterson, and R. C. Radin. “Two Pretreatment Regimens for High-Risk Patients Receiving Radiographic Contrast Media.” J Allergy Clin Immunol 74, no. 4 Pt 1 (1984): 540–3.

57. Gabizon, A., R. Catane, B. Uziely, B. Kaufman, T. Safra, R. Cohen, F. Martin, A. Huang, and Y. Barenholz. “Prolonged Circulation Time and Enhanced Accumulation in Malignant Exudates of Doxorubicin Encapsulated in Polyethylene-Glycol Coated Liposomes.” Cancer Res 54, no. 4 (1994): 987–92.

58. Gabizon AA, Gabizon-Peretz S, Modaresahmadi S, La-Beck NM. Thirty years from FDA approval of pegylated liposomal doxorubicin (Doxil/Caelyx): an updated analysis and future perspective. BMJ Oncol. 2025;4(1):e000573.

59. Gabizon, A., and F. Martin. “Polyethylene Glycol-Coated (Pegylated) Liposomal Doxorubicin. Rationale for Use in Solid Tumours.” Drugs 54 Suppl 4 (1997): 15–21.

60. Gabizon, A. A. “Pegylated Liposomal Doxorubicin: Metamorphosis of an Old Drug into a New Form of Chemotherapy.” Cancer Invest 19, no. 4 (2001): 424–36.

61. Botting, R. M. “Mechanism of Action of Acetaminophen: Is There a Cyclooxygenase 3?” Clin Infect Dis 31 Suppl 5 (2000): S202–10.

62. Szebeni, Janos, John L Fontana, Nabila M Wassef, Paul D Mongan, David S Morse, David E Dobbins, Gregory L Stahl, Rolf Bunger, and Carl R Alving. “Hemodynamic Changes Induced by Liposomes and Liposome-Encapsulated Hemoglobin in Pigs: A Model for Pseudoallergic Cardiopulmonary Reactions to Liposomes: Role of Complement and Inhibition by Soluble Cr1 and Anti-C5a Antibody.” Circulation 99, no. 17 (1999): 2302–09.

63. Urbanics, R., and J. Szebeni. “Lessons Learned from the Porcine Carpa Model: Constant and Variable Responses to Different Nanomedicines and Administration Protocols.” Eur J Nanomedicine 7 (2015): 219–31.

64. Szebeni, J., and R. Bawa. “Human Clinical Relevance of the Porcine Model of Pseudoallergic Infusion Reactions.” Biomedicines 8, no. 4 (2020).

65. Szebeni, J. “Evaluation of the Acute Anaphylactoid Reactogenicity of Nanoparticle-Containing Medicines and Vaccines Using the Porcine CARPA Model.” In Characterization of Nanoparticles Intended for Drug Delivery, 10.1007/978-1-0716-3786-9_23,, edited by Jeffrey D. Clogston et al. (eds.). 10.1007/978-1-0716-3786-9_23, 2024.

66 Barroso, A., F. Estevinho, V. Hespanhol, E. Teixeira, J. Ramalho-Carvalho, and A. Araújo. “Management of Infusion-Related Reactions in Cancer Therapy: Strategies and Challenges.” ESMO Open 9, no. 3 (2024).66. 10.1016/j.esmoop.2024.102922.

